# Analysis of the small chromosomal *Prionium serratum* (Cyperid) demonstrates the importance of a reliable method to differentiate between mono- and holocentricity

**DOI:** 10.1101/2020.07.08.193714

**Authors:** M. Baez, Y.T. Kuo, Y. Dias, T. Souza, A. Boudichevskaia, J. Fuchs, V. Schubert, A.L.L. Vanzela, A. Pedrosa-Harand, A. Houben

**Affiliations:** Leibniz Institute of Plant Genetics and Crop Plant Research (IPK) Gatersleben, 06466 Stadt Seeland, Germany; Laboratory of Plant Cytogenetics and Evolution, Department of Botany, Federal University of Pernambuco, Pernambuco, Brazil; Laboratory of Cytogenetics and Plant Diversity, Department of General Biology, Center for Biological Sciences, State University of Londrina, Londrina 86057-970, Paraná, Brazil; KWS SAAT SE & Co. KGaA, 37574, Einbeck, Germany

**Keywords:** CENH3/CENPA, centromere type, holocentric chromosome, evolution, Cyperids, Thurniceae

## Abstract

For a long time, the Cyperid clade (Thurniceae-Juncaceae-Cyperaceae) was considered as a group of species possessing holocentromeres exclusively. The basal phylogenetic position of *Prionium serratum* L. f. Drège (Thurniceae) within Cyperids makes this species an important specimen to understand the centromere evolution within this clade. Unlike expected, the chromosomal distribution of the centromere-specific histone H3 (CENH3), alpha-tubulin and different centromere associated post-translational histone modifications (H3S10ph, H3S28ph and H2AT120ph) demonstrate a monocentromeric organisation of *P. serratum* chromosomes. Analysis of the high-copy repeat composition resulted in the identification of a centromere-localised satellite repeat. Hence, monocentricity was the ancestral condition for the Juncaceae-Cyperaceae-Thurniaceae Cyperid clade and holocentricity in this clade has independently arisen at least twice after differentiation of the three families, once in Juncaceae and the other one in Cyperaceae. Methods suitable for the identification of holocentromeres are discussed.

## Introduction

Centromeres are essential for the segregation of chromosomes to the daughter cells during mitosis and meiosis. Most organisms contain one single size-restricted centromere per chromosome (monocentromere) visible as a primary constriction during metaphase. However, in independent eukaryotic taxa, species with chromosomes without distinct primary constrictions visible at metaphase exist, which are referred to as holocentric. Instead, the spindle fibres attach along almost the entire poleward surface of the chromatids (reviewed in Schubert *et al.* (2020)). Holocentricity evolved at least 19 times independently in various green algae, protozoans, invertebrates, and different higher plant families (Dernburg, 2001; Escudero *et al.*, 2016; Melters *et al.*, 2012). In total, ~1.5 - 2.0% of flowering plants are likely to have holocentric chromosomes (Bures *et al.*, 2012). It is possible that holocentricity is even more common than reported so far, as the identification of the centromere type, especially in small-sized chromosomes, is challenging.

One common explanation for the evolution of holocentric chromosomes is their putative advantage related to DNA double-strand breaks (Zedek and Bures, 2018). The studies on artificial chromosomal rearrangements in various holocentric species showed that chromosome fragments retaining centromere activity are transmitted during mitosis and meiosis (Jankowska *et al.*, 2015). Comparisons of diversification rates between monocentric and holocentric sister clades in animals and plants did not detect an increase in diversification in holocentric species (Marquez-Corro *et al.*, 2018). Nevertheless, these analyses depend on the correct identification of the centromere type in a large number of lineages.

Because holocentric taxa are often embedded within broader phylogenetic lineages possessing monocentric chromosomes, it is thought that holocentric chromosome organisation originated from the monocentrics and that this transition occurred independently in multiple phylogenetic lineages (Melters *et al.*, 2012). However, the factors that induced this transition and its mechanisms are currently unknown. Investigations of the changes associated with the transition from monocentric to holocentric chromosome organisation are, in theory, most informative when phylogenetically closely related species that differ in the centromere type are compared.

In angiosperms, holocentric chromosomes have been confirmed in some dicot species, e.g. in the genus *Cuscuta* L. subgenus *Cuscuta* (Convolvulaceae) (Oliveira *et al.*, 2020) and in a few species within the genus *Drosera* L. (Droseraceae) (Sheikh *et al.*, 1995). Also, in monocots, for example, in the genus *Luzula* DC (Juncaceae) (Heckmann *et al.*, 2013) and *Rhynchospora* Vahl. (Cyperaceae) (Marques *et al.*, 2015; Ribeiro *et al.*, 2017) holocentricity occurs. These last two families belong to the Cyperid clade (Thurniceae-Juncaceae-Cyperaceae), which was originally considered to share holocentric chromosomes as a synapomorphic feature (Greilhuber, 1995; Judd *et al.*, 2016; Melters *et al.*, 2012). However, exceptions have been reported in the genus *Juncus* L., in which four species exhibited primary constrictions (Marcelo Guerra *et al.*, 2019). It suggests that this synapomorphy of the Cyperid clade is uncertain.

Aiming to improve the understanding on the origin and evolution of the holocentricity within the Cyperid clade we studied the centromere organisation of *Prionium serratum* L. f. Drège (Thurniceae), a species phylogenetically situated at the base of the Cyperid clade (Silva *et al.*, 2020) (Supp. Figure 1). The South African monocotyledonous plant genus *Prionium* E. Mey is an old, species-poor lineage which split from its sister genus about 26.1 million years ago (Kumar *et al.*, 2017). *P. serratum* is suspected to be holocentric, as it is closely related to the families Juncaceae and Cyperaceae. Supported was this assumption by the fact that this species has a low genomic GC content, as it is typically described for holocentric species (Smarda *et al.*, 2014). Furthermore, Zedek et al. (2016) observed no significant increase in the proportion of G2 nuclei after gamma irradiation of *P. serratum,* differing from monocentric species.

To ascertain the centromere type of *P. serratum,* we determined the chromosomal distribution of the centromere-specific histone H3 (CENH3) protein, alpha-tubulin, histone H3 phosphorylated at position serine 10, serine 28 (H3S10ph, H3S28ph) and histone H2A phosphorylated at position threonine 120 (H2AT120ph). The cell cycle-dependent post-translational histone modifications are associated with active centromeres (Demidov *et al.*, 2014; Dong and Han, 2012; Gernand *et al.*, 2003). Unlike expected, a monocentromeric organisation of the chromosomes was found. The analysis of the high-copy repeat composition resulted in the identification of two centromere-localised satellite repeats. In addition, a DNA replication behaviour was found typical for small genome monocentric species. The data are discussed in the context of centromere evolution in Cyperids and concerning the suitability of available methods to identify holocentromeres.

## Materials and methods

### Plant material

Individuals of *Prionium serratum* L. f. Drège, collected in western Cape (Cape Town, South Africa; TE2016_413) and provided by the Herrenhäuser Gardens (Hannover, Germany, IPK herbarium 70142) and the Botanical Garden Halle (Halle, Germany) were grown in a greenhouse of the Leibniz Institute of Plant Genetics and Crop Plant Research (IPK Gatersleben, Germany).

### Flow cytometric genome size measurement

For nuclei isolation, roughly 0.5 cm^2^ of fresh leaf tissue were chopped together with equivalent amounts of leaf tissue of one of the internal reference standards, *Raphanus sativus* var. ‘Voran’ (Gatersleben genebank accession number: RA 34; 1.11 pg/2C) or *Lycopersicon esculentum* var. ‘Stupicke Rane’ (Gatersleben genebank accession number: LYC 418; 1.96 pg/2C), in a petri dish using the reagent kit ‘CyStain PI Absolute P’ (Sysmex) following the manufacturer’s instructions. The resulting nuclei suspension was filtered through a 50 μm filter mesh (CellTrics, Sysmex) and measured either on a CyFlow Space (Sysmex) or on a BD Influx cell sorter (BD Biosciences). The absolute DNA content (pg/2C) was calculated based on the values of the G1 peak means and the corresponding genome size (Mbp/1C), according to Dolêzel et al. (2003).

### DNA/RNA extraction and sequencing

Genomic DNA was extracted from *P. serratum* leaves using the DNeasy Plant Mini Kit (Qiagen) and sequenced using the HiSeq 2500 system (Illumina, CA) at low coverage. RNA was extracted from root meristems and prepared for paired-end sequencing on Illumina HiSeqX (Illumina, CA) by Novogene (Beijing, China).

### In silico repeat analysis

The repetitive proportion of the genome was analysed by the RepeatExplorer pipeline (Novak *et al.*, 2013), implemented within the Galaxy/Elixir environment (https://repeatexplorer-elixir.cerit-sc.cz/). Low-coverage genomic paired reads were filtered by quality with 95% of bases equal to or above the quality cut-off value of 10 and interlaced. Clustering was performed with a minimum overlap of 55% and a similarity of 90%. Protein domains were identified using the tool Find RT Domains in RepeatExplorer pipeline (Novak et al., 2013). Searches using databases (GenBank) were performed and graph layouts of individual clusters were examined interactively using the SeqGrapheR program (Novak *et al.*, 2013).

The number of analysed reads was 2,752,532 comprising in total ~276 Mbp, corresponding to 0.82x genome coverage. All clusters representing at least 0.01% of the genome were manually checked, and their automated annotation was corrected if necessary. The size of the annotated clusters was used to characterise and quantify the genome proportion of the high-copy repeats. To reconstruct the conserved monomer sequence of the tandem repeats, three independent runs were performed using the TAREAN (TAndem REpeat ANalyzer) tool implanted in RepeatExplorer (Novak et al., 2017).

### Repeat amplification, probe labelling and fluorescent in situ hybridisation

Satellite DNAs (satDNA) were PCR amplified with primers facing outwards of a repeat unit or directly synthesised as oligonucleotides with 5’-labelled fluorescence. Primers and oligonucleotides were designed from the most conserved region of the consensus sequences (Table 1). Forty nanograms of genomic DNA were used for all PCR reactions with 1× PCR buffer, 2 mM MgCl_2_, 0.1 mM of each dNTP, 0.4 μM of each primer, 0.025 U *Taq* polymerase (Qiagen) and water. PCR condition was: 94°C 3 min, 30× (94°C 1 min, 55°C 1 min, 72°C 1 min) and 72°C 10 min. Amplicons and plasmid DNA of the 45S rDNA-containing clone pTa71 (Gerlach and Bedbrook, 1979) were labelled with either Cy3, Atto488 or **Atto550 fluorophores** by a nick translation labelling kit (Jena Bioscience).

**Table 1.**
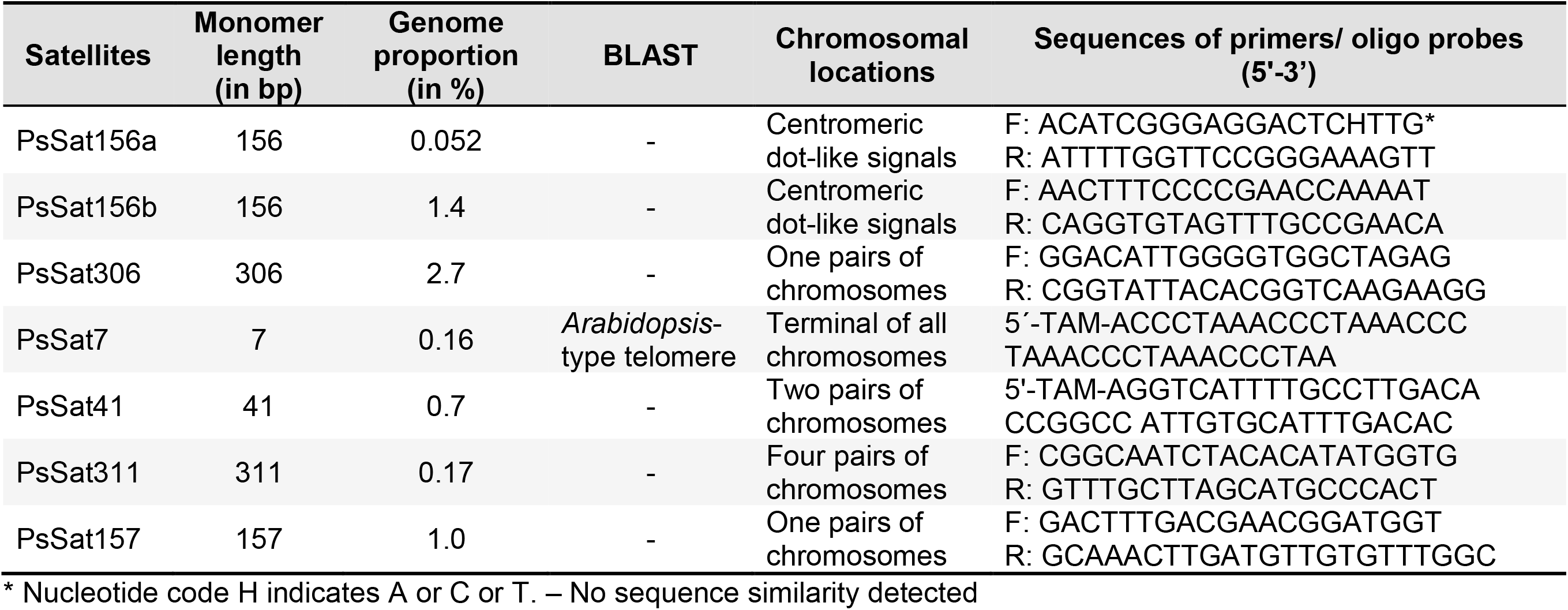
High-copy satellite repeats of *P. serratum* and their corresponding chromosomal localisations

Mitotic chromosomes were prepared from root tips, pre-treated in 2 mM 8-hydroxyquinoline at 7°C for 24 h and fixed in ethanol: acetic acid (3:1 v/v) for 2 to 24 h at room temperature and stored at −20°C. Fixed root tips were digested with 2% cellulose, 2% pectinase, 2% pectolyase in citrate buffer (0.01 M sodium citrate dihydrate and 0.01 M citric acid) for 90 min at 37°C and squashed in a drop of 45% acetic acid. Fluorescent *in situ* hybridisation was performed as described by Aliyeva-Schnorr et al. (2015). The hybridisation mix contained 50% (v/v) formamide, 10% (w/v) dextran sulfate, 2× SSC, and 5 ng/μl of each probe. Slides were denatured at 75°C for 5 min, and the final stringency of hybridisation was 76%.

### RNA sequence analysis

We generated a total 15.6 Gbp of paired-end reads of 150 bp (around 52 million reads per end). Prior to mapping, all reads were preprocessed for quality control with FastQC, Galaxy version 0.72 (Andrews, 2010). Subsequently, they were processed with the Trimmomatic program, Galaxy version 0.36.6 (Bolger *et al.*, 2014) to trim adaptor contamination and low-quality sequences. As a result, 93.9% of high‐quality sequences from total number were used for *de novo* transcriptome assembly with Trinity version 2.4.0. To evaluate the quality of assembly, its completeness and to remove poorly supported contigs, we applied Transrate v1.0.3 (Smith-Unna *et al.*, 2016). The resulting dataset with 68,922 contigs was further processed by CD-HIT-EST, v. 4.6.8 program, using -c 0.95 -n 10 as parameters (Fu *et al.*, 2012; Li and Godzik, 2006) to cluster highly-homologous sequences and remove redundant transcripts. Afterwards, the resulting file with 67,565 contigs was used to identify candidate coding regions within the transcript sequences (Transdecoder v. 5.3.0; http://transdecoder.github.io). RNAseq data are deposited in the European Nucleotide Archive under PRJEB39221, genomic data are under NCBI SRX8683442. To identify a CENH3 candidate in the RNA-seq data we performed BLASTP, Galaxy Version 0.3.3 (Cock et al. 2015) using CENH3s from other monocotyledonous plants.

### Phylogenetic analysis

The CENH3 sequence selected from *P. serratum* transcriptome dataset and those of other species downloaded from NCBI GenBank (see Figure 1) were aligned with ClustalW implanted in MEGA X, using the default setting (Kumar *et al.*, 2018; Thompson *et al.*, 1994). The evolutionary relationship was inferred using the maximum likelihood method by IQ-Tree web server (http://iqtree.cibiv.univie.ac.at) (Trifinopoulos *et al.*, 2016). The built tree was visualised, labelled and exported by Interactive Tree Of Life (iTOL, https://itol.embl.de/) (Letunic and Bork, 2007, 2019).

**Figure 1.**
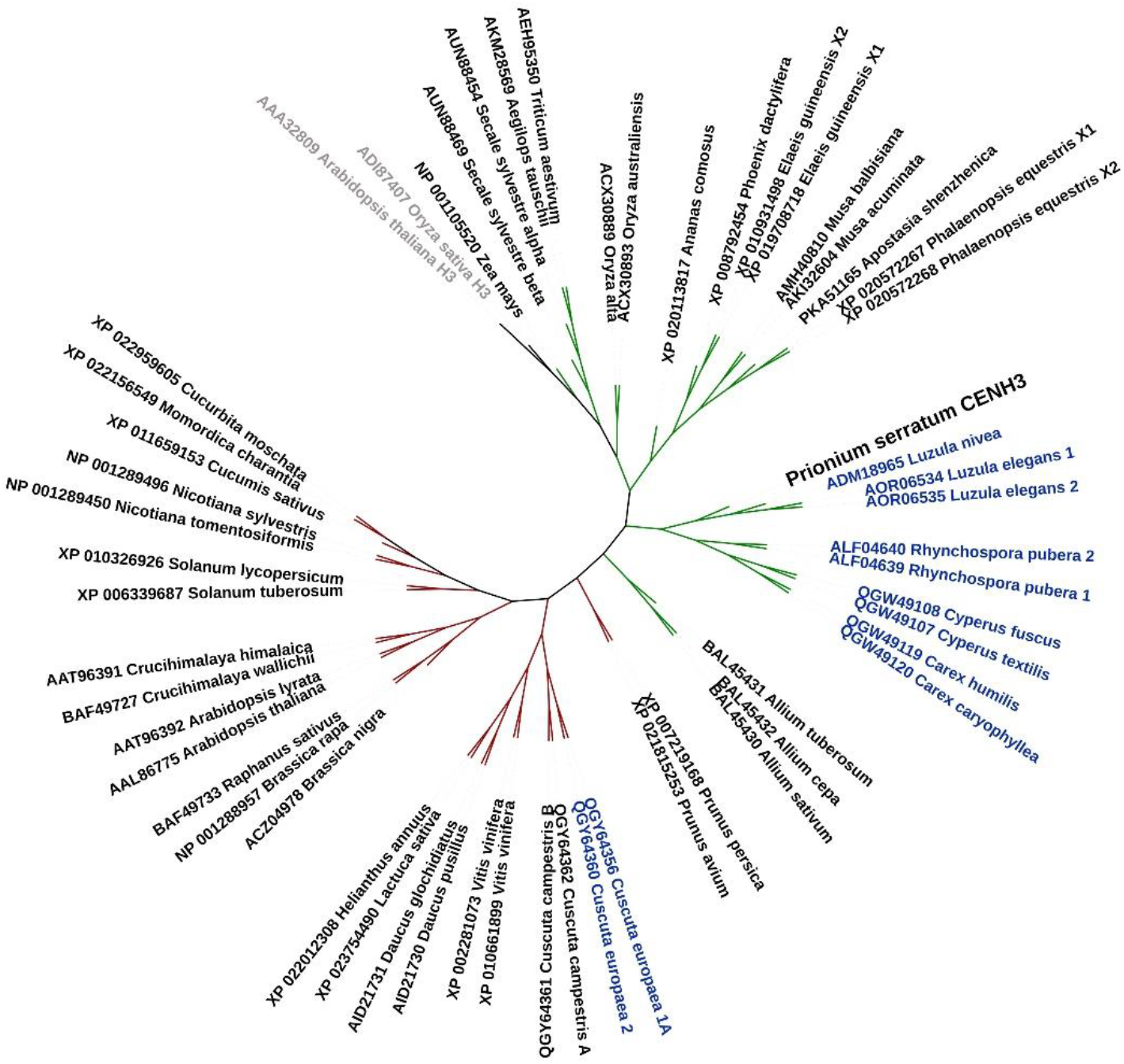
Phylogenetic relationships of CENH3 between *P. serratum* and other plant species. The green and red branch represent monocot and eudicot species, respectively. The blue node indicates the reported holocentric species and the sequences of the canonical histone 3 (H3) used as outgroup are shown in grey node. The accession number of the CENH3 derived from the NCBI GenBank is included in the node label.

### Indirect immunostaining

The PsCENH3: RVKHFSNKAVSRTKKRIGSTR-c peptide was used for the production of polyclonal antibodies in rabbits. LifeTein (www.lifetein.com) performed the peptide synthesis, immunisation of rabbits, and peptide affinity purification of antisera. Mitotic preparations were made from root meristems fixed in paraformaldehyde and Tris buffer (10 mM Tris, 10 mM EDTA, 100 mM NaCl, 0.1% Triton, pH 7.5) or 1×MTSB buffer (50 mM PIPES, 5 mM MgSO_4_, and 5 mM EGTA, pH 7.2) for 5 minutes on ice in a vacuum and for another 25 minutes only on ice. After washing twice in Tris buffer or 1×MTSB buffer, the roots were chopped in LB01 lysis buffer (15 mM Tris, 2 mM Na_2_EDTA, 0.5 mM spermine∙4HCl, 80 mM KCl, 20 mM NaCl, 15 mM β-mercaptoethanol, 0.1% (v/v) Triton X-100, pH 7.5), filtered through a 50 μm filter (CellTrics, Sysmex) and diluted 1:10, and subsequently, 100 μl of the diluted suspension were centrifuged onto microscopic slides using a Cytospin3 (Shandon, Germany) as described (Jasencakova *et al.*, 2001). Immunostaining was performed as described by Houben *et al.* (2007). The following primary antibodies were used: rabbit anti-PsCENH3 (diluted 1:300), mouse anti-alpha-tubulin (clone DM 1A, Sigma, diluted 1:200), mouse anti-histone H3S10ph (Abcam, 14966, diluted 1:200), mouse anti-histone H3S28ph (Millipore, 09_797, diluted 1:200) and rabbit anti-histone H2A120ph ((Demidov *et al.*, 2014), diluted 1:200). As secondary antibodies, a Cy3-conjugated anti-rabbit IgG (Dianova) and a FITC-conjugated anti-mouse Alexa488 antibody (Molecular Probes) were used in a 1:500 dilution each. Slides were incubated overnight at 4 °C, washed 3 times in 1×PBS or 1×MTSB and then the secondary antibodies were applied. Immuno-FISH was performed, according to Ishii *et al.* (2015).

### DNA replication analysis

Roots were treated for 2 h with 15 μM EdU (5-ethynyl-2’-deoxyuridine, baseclick GmbH), followed by water for 30 minutes. Preparation of slides was performed as described for immunostaining. The click reaction was performed to detect EdU according to the manual (baseclick GmbH).

### Microscopy

Images were captured using an epifluorescence microscope BX61 (Olympus) equipped with a cooled CCD camera (Orca ER, Hamamatsu). To achieve super-resolution of ~120 nm (with a 488 nm laser excitation), we applied spatial structured illumination microscopy (3D-SIM) using a 63x/1.40 Oil Plan-Apochromat objective of an Elyra PS.1 microscope system (Carl Zeiss GmbH) (Weisshart *et al.*, 2016).

## Results

### *Prionium serratum* is a monocentric species

*Prionium serratum* was chosen to test whether holocentricity occurs at the base of the Cyperid clade, since this species is phylogenetically positioned at the base of a group of species recognised as holocentrics. Since the around 1 μm long mitotic metaphase chromosomes did not allow the doubtless identification of a monocentromere-typical primary constriction or a holocentromere-typical parallel configuration of anaphase sister chromatids, we generated a CENH3-specific antibody suitable for immunostaining. The centromere-specific histone variant CENH3 was shown to be essential for centromere function in many species (Allshire and Karpen, 2008).

First, the root transcriptome of *P. serratum* was determined and the assembled RNAseq reads were used to identify *CENH3.* Only one *CENH3* gene was identified in the transcriptome dataset. After alignment of the corresponding amino acid sequence against CENH3s of other plant species, the evolutionary tree grouped *P. serratum* CENH3 together with other Cyperid sequences belonging *to Luzula* (Juncaceae), *Rhynchospora*, *Cyperus*, and *Carex* (Cyperaceae), supporting the correct identification of the CENH3 gene (Figure 1).

Next, antibodies (anti-PsCENH3) designed to recognise CENH3 of *P. serratum* were generated and used for immunostaining. Typical monocentromere dot-like signals were found at interphase and at early prophase (Figure 2A, B). The observed interaction of CENH3 with alpha-tubulin fibres at metaphase proved the centromere specificity of the CENH3 signals (Figure 2C, D). Besides, the cell-cycle dependent, pericentromere-specific distribution of H3S10ph, H3S28ph and H2AT120ph confirmed a monocentric chromosome type (Figure 3A-C). Hence, *P. serratum* is monocentric species based on the results obtained by the application of different (peri)centromere-specific antibodies.

**Figure 2.**
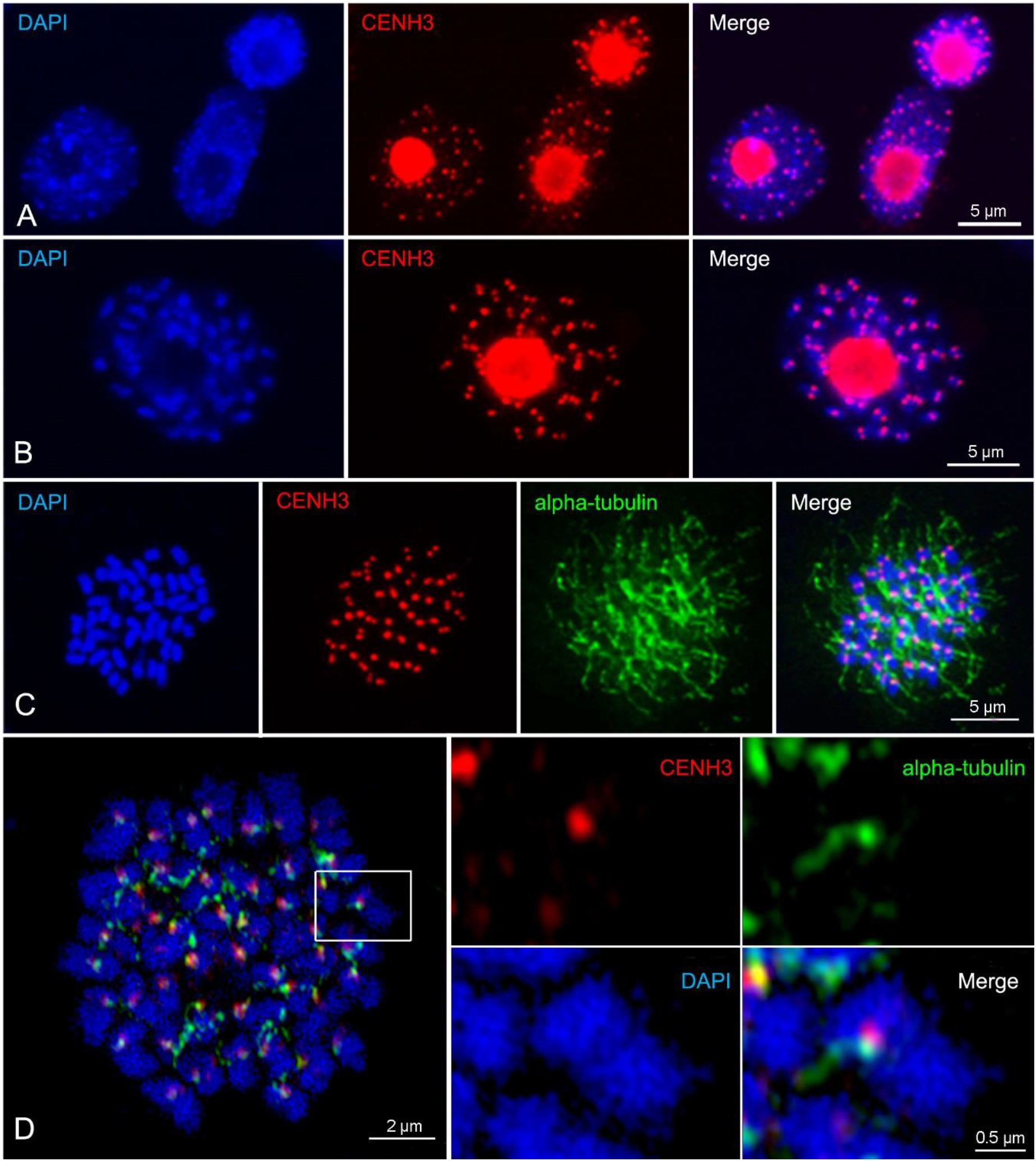
Immunodetection of centromeric protein CENH3 (red) in *P. serratum* interphase nuclei (A), prophase (B) and its interaction with alpha-tubulin (green) in metaphase chromosomes (C, D). (D) Image taken by spatial structured illumination microscopy (SIM), enlargement (square) shows the interaction between CENH3 and alpha-tubulin.

**Figure 3.**
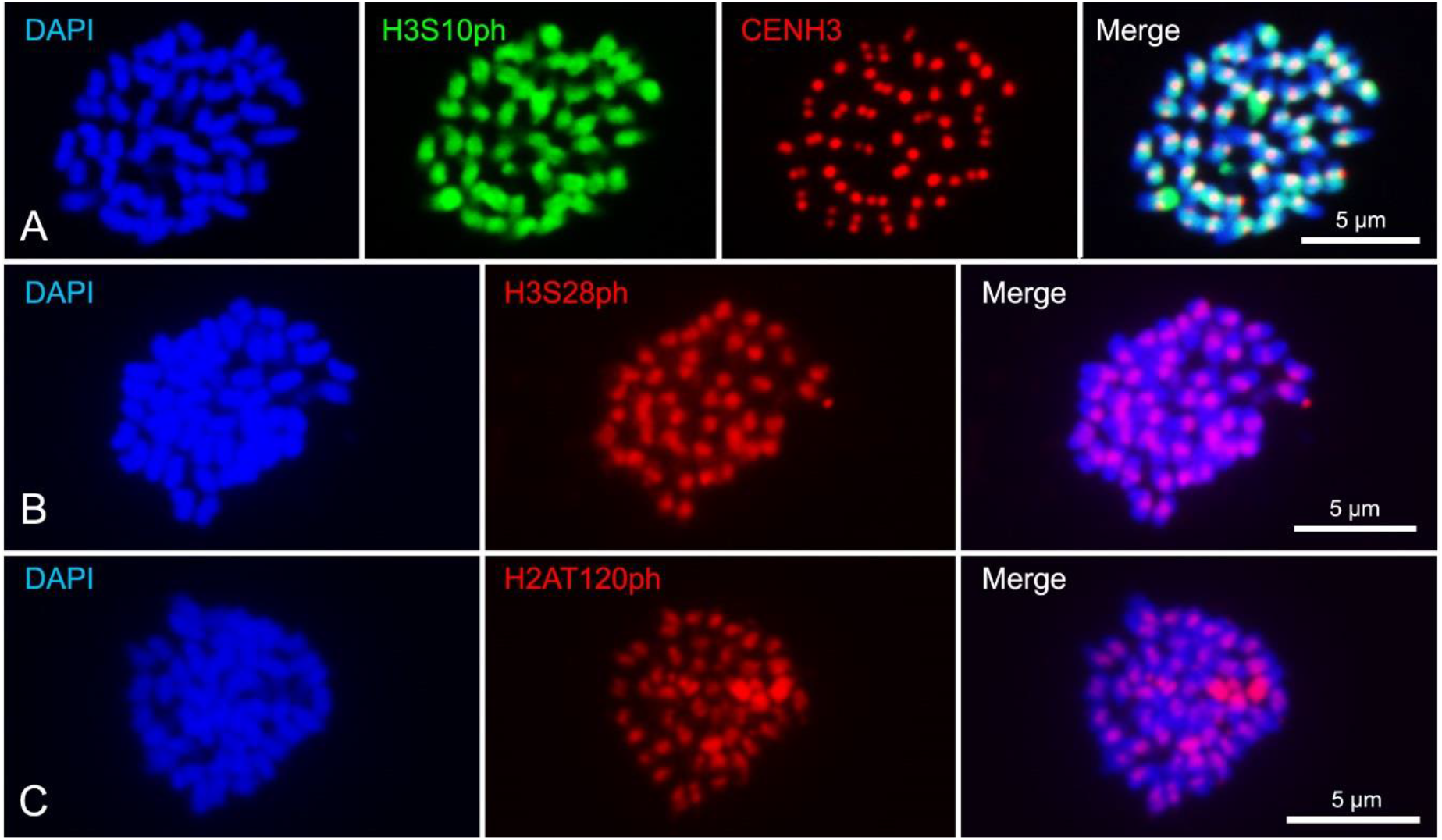
Cell-cycle dependent, pericentromere-specific histone phosphorylated modification at H3S10 (A), H3S28 (B) and H2AT120 (C) in metaphase chromosomes of *P. serratum*. Overlapped signals between H3S10ph (green) and CENH3 (red) are shown in (A).

### Identification of a centromere-localised repeat family *in P. serratum*

The genome size of *P. serratum* (2*n*=46) is 335 Mbp/1C, estimated by flow-cytometry. Next-generation sequence reads were generated to investigate the repetitive composition of the *P. serratum* genome based on the graph-based clustering analysis, resulting in the identification of high-copy satellite repeats and transposable elements. About 26.9% of the genome is composed of repetitive elements. The top first 329 clusters with at least 0.01% genome proportion, classified as 13 lineages of class I transposable elements (LTR-retrotransposons and non-LTR LINE), six class II DNA transposons, satellite DNA (satDNA) as well as ribosomal DNA (rDNA) (Table 2). The LTR-retrotransposons constituted ~9% of the genome, with the Ty1-Copia elements being more abundant than the Ty3-Gypsy elements, representing genome proportions of 5.36% and 3.63%, respectively.

**Table 2.**
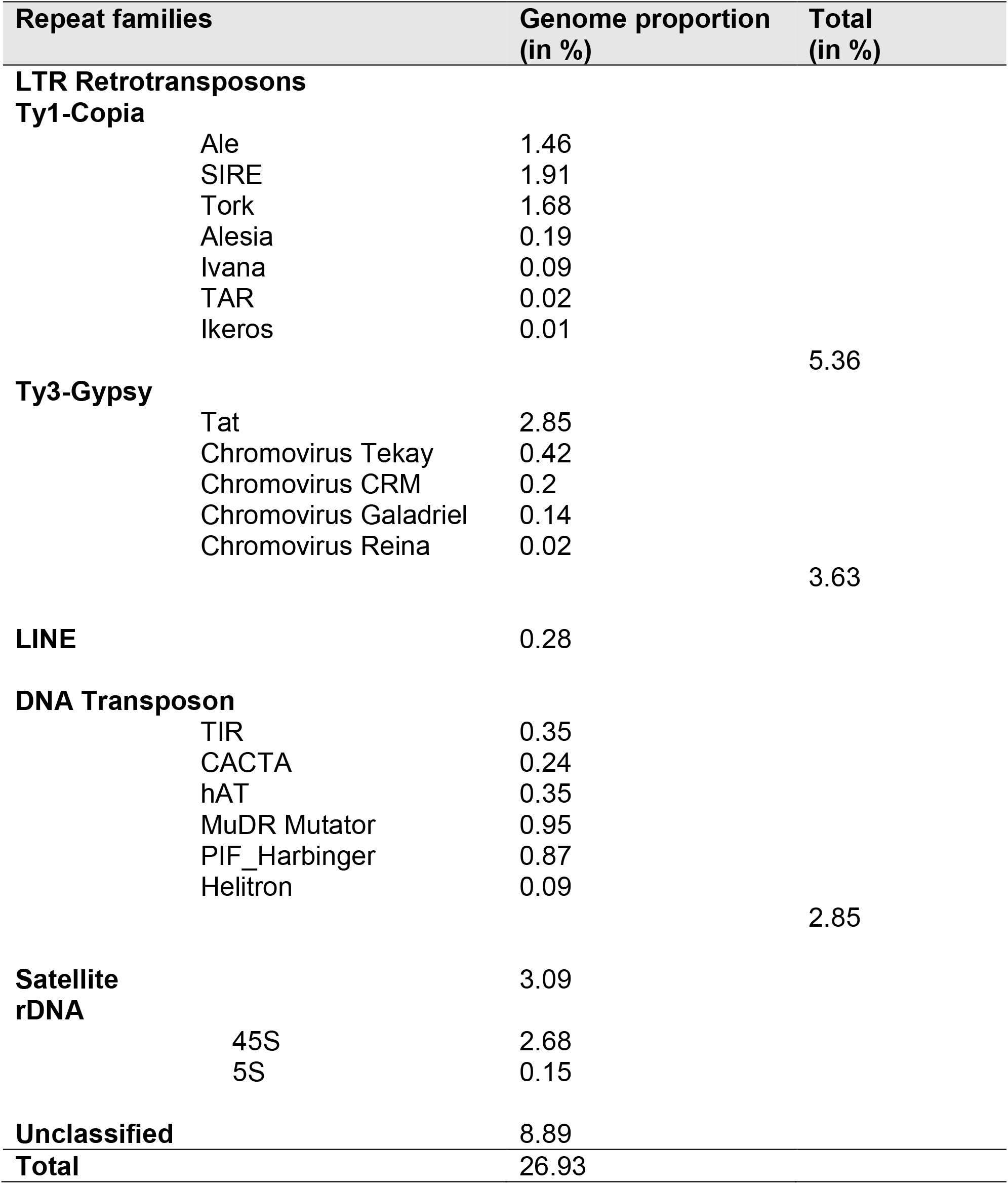
Repetitive families of *P. serratum*

The k-mer-based TAREAN analysis resulted in the identification of 19 different satDNA families. Out of these, the seven most abundant satDNAs were used for FISH to determine their chromosomal distribution. PsSat7, representing the *Arabidopsis*-type telomere sequence, hybridised to the terminal regions of all chromosomes. It is likely that the copy number of telomere repeats differs between the individual chromosome ends, as the intensity of the signals varied (Figure 4A). PsSat41, PsSat311 and PsSat157clustered on two, four and one chromosome pairs, respectively (Figure 4B, C, D). PsSat306 colocalised with 45S rDNA signals (Figure 4E).

**Figure 4.**
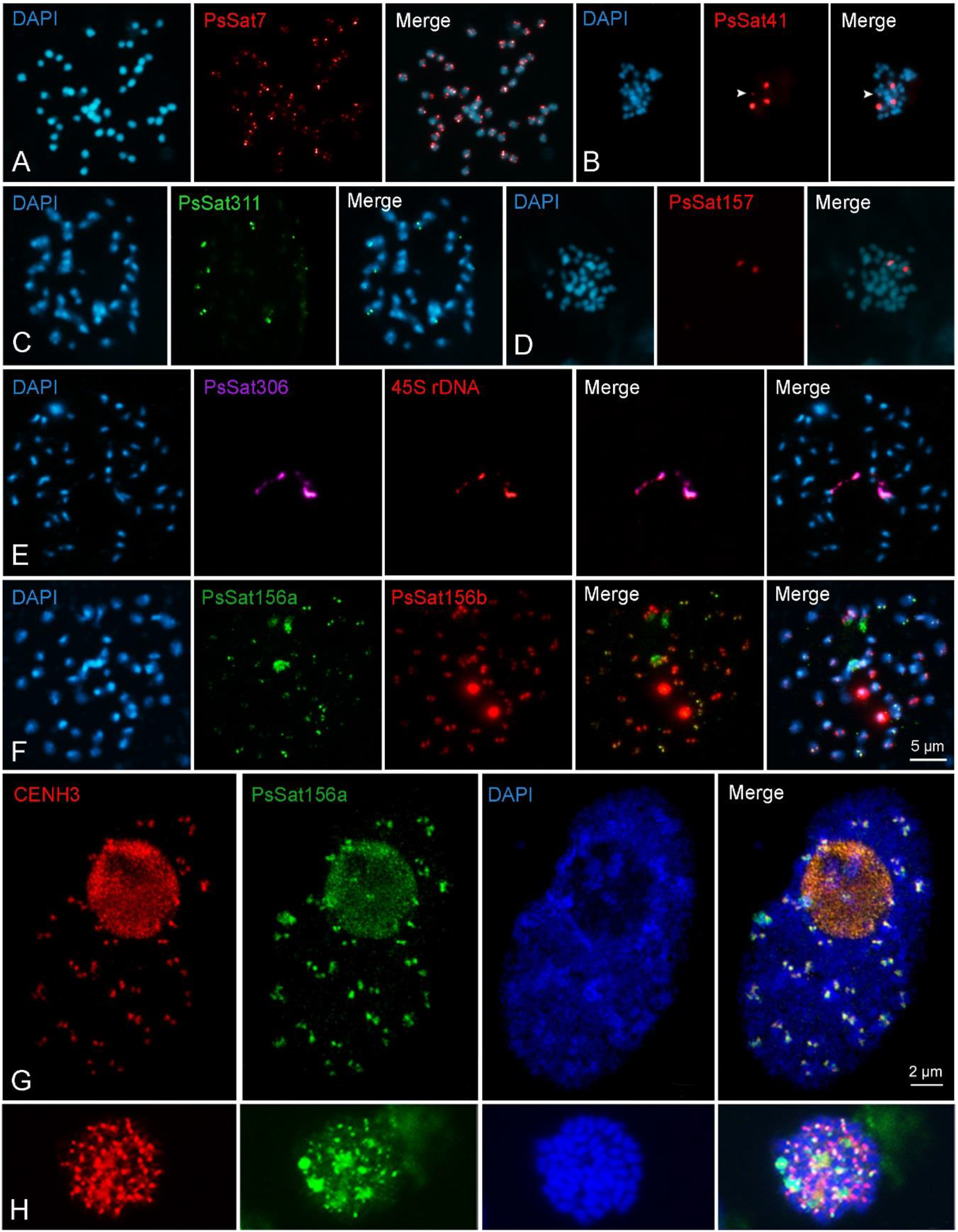
Chromosome distribution of satellite DNA families and of CENH3 in *P. serratum* (2*n* = 46). Satellite repeat PsSat7 (A), PsSat41 (B), PsSat311 (C), PsSat157 (D), PsSat306 (E), Ps156a and Ps156b (F) were mapped on metaphase chromosomes. The fourth signal of PsSat41 is indicated by *arrowheads* (B). Colocalisation between PsSat306 and 45S rDNA and between Ps156a and Ps156b are shown in (E) and (F), respectively. The centromere specificity of Ps156a was confirmed by its overlapped signals, visualised in *yellow* in the merge images, with CENH3 in both interphase nuclei (G) and metaphase chromosomes (H). Image (G) was taken by structured illumination microscopy (SIM).

Centromere-like signals were only found after FISH with the satDNA family PsSat156 (Figure 4F). PsSat156a and PsSat156b possess a sequence similarity of 96% but with different abundance at chromosomes. Beside dot-like signals, both probes showed enlarged hybridisation signals on one but different chromosome pairs each. To confirm the centromeric position of PsSat156, immuno-FISH with the CENH3-specific antibody was performed. Colocalisation of both signals in metaphase chromosomes and interphase nuclei demonstrated the centromere specificity of the repeat family PsSat156 (Figure 4G). No sequence similarity was found between PsSat156 and centromeric repeats of other species.

Finally, the DNA replication behaviour was studied to test whether early‐ and late‐replicating chromosome regions occupy distinct chromosomal regions, as in most monocentric species (Costas *et al.*, 2011). 5-ethynyl-2’-deoxyuridine (EdU), a nucleoside analogue of thymidine, was applied during DNA replication of *P. serratum* and two major types of nuclei labelling patterns were found (Figure 5). The majority of nuclei (85% of 500 nuclei) showed an almost uniform labelling (Figure 5A) and 15% of nuclei showed a cluster-like distribution of EdU signals (Figure 5B). Comparable replication patterns were found in other species with monocentric chromosomes like *Arabidopsis thaliana* (Dvorackova *et al.*, 2018) and *Zea mays* (Bass *et al.*, 2015).

**Figure 5.**
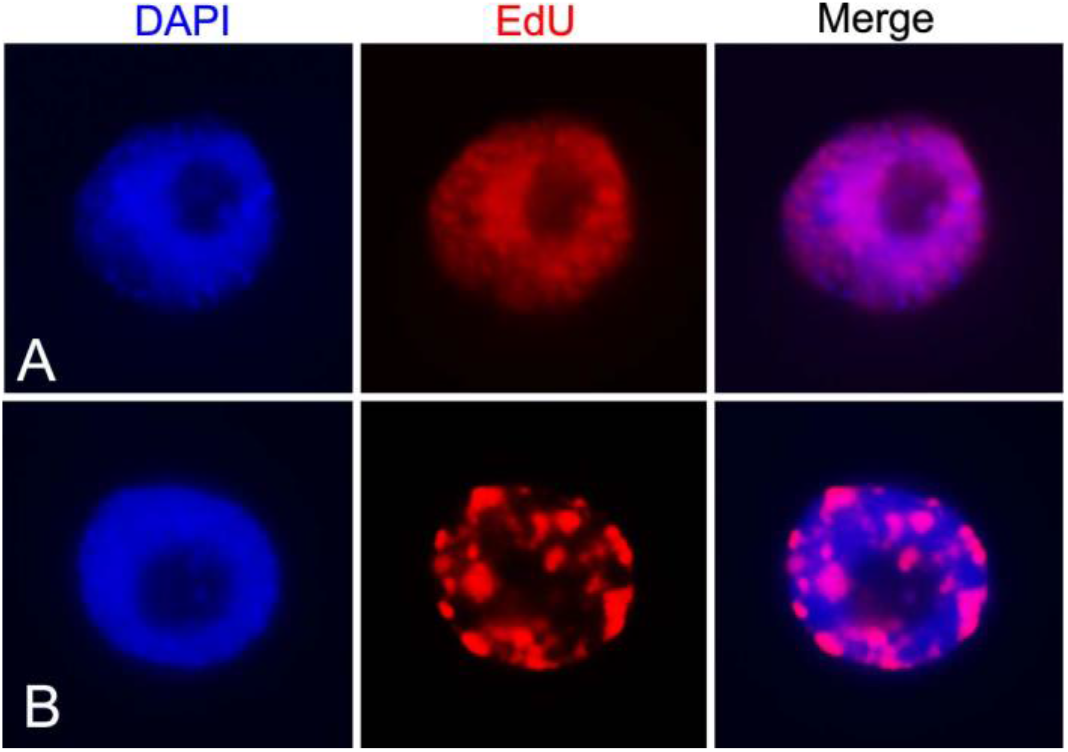
Two types of DNA replication patterns in *P. serratum* shown by EdU labelling (red) and interphase nuclei counterstained with DAPI (blue). (A) Mainly uniform labelling and (B) clustered distribution of EdU signals.

## Discussion

### Centromere evolution in Cyperids

The analysis of the centromeres by immunostaining using CENH3, alpha-tubulin, histone H3S10ph, H3S28ph and H2AT120ph antibodies demonstrated a monocentric centromere-type for the phylogenetically basal *P. serratum.* Therefore, these data suggest that monocentric chromosomes may be an ancestral condition for the Juncaceae-Cyperaceae-Thurniaceae Cyperid clade. As monocentricity was also reported in species within the *Juncus* genus (Guerra *et al.*, 2019), holocentric chromosomes in the Cyperid clade have evolved at least twice independently: once in Juncaceae and once in Cyperaceae, after the divergence of the three families. The phylogenetic close proximity of *P. serratum* CENH3 with species possessing holocentromere demonstrates that the sequence divergence of CENH3 does not correlate with its corresponding centromere type. A similar centromere-type independent CENH3 evolution was found for mono- and holocentric *Cuscuta* species (Oliveira *et al.*, 2020). Hence, our data suggested that sequence modifications of CENH3 are not necessarily involved in the change of the centromere-type in angiosperms.

### The abundance of repetitive DNA in *P. serratum*

The small *P. serratum* genome contains a relatively low percentage of transposable elements, ~9% retrotransposons and ~3% transposons sequences, with 10 and 6 different lineages, respectively. Most plant genomes contain only a few satDNA families, mainly repeats associated with pericentromeric or subtelomeric regions (reviewed in (Garrido-Ramos, 2015). Here, we identified 19 different satDNA families (~3% of the genome). Unlike in other small genome sized, monocentric species, like *A. thaliana* (Maluszynska and Heslop Harrison, 1991), sugar beet (Kubis *et al.*, 1998) and rice (Cheng *et al.*, 2002), the centromeric satDNA is not the most abundant satDNA. PsSat306, the most abundant satDNA family, displays colocalisation with the 45S rDNA. PsSat306 likely originated from the intergenic repeat spacer region as described for other satellite repeats (reviewed in (Garrido-Ramos, 2015)). In addition, the 5-ethynyl-2’-deoxyuridine (EdU) detection in interphase showed comparable late replication patterns those observed in species with monocentric chromosomes (Bass *et al.*, 2015; Dvorackova *et al.*, 2018).In holocentric species like *L. elegans,* the chromosomes are less clearly compartmentalised into distinguishable early‐ and late‐replicating chromosome regions (Heckmann *et al.*, 2013).

A cluster distribution at one or only a few chromosome pairs was found for three satDNA families, similar to others satellite repeats in several species within the clade, as the holocentric *Luzula* and *Rhynchospora* genera, and outside the clade in typical monocentric species, as *Chenopodium quinoa* (Heckmann *et al.*, 2013; Heitkam *et al.*, 2020; Ribeiro *et al.*, 2017). Two of these tandem repeats (PsSat156a and PsSat156b) share the same distribution at centromeric regions but with different signal intensities. Most likely, they evolved from the same ancestral centromeric repeat unit, and underwent amplification or reduction at different chromosome pairs.

### How to identify holocentricity?

Results in *P. serratum* demonstrated that the characterisation of the centromere type, especially in species with small-sized chromosomes could be challenging. Which is the best method to identify holocentricity? As listed in Table 3, a range of different methods has been used to determine the centromere type in the past. However, no universal method amenable for all species exists, either due to the limitation in optical resolution, availability of specific antibodies or required equipment.

**Table 3.**
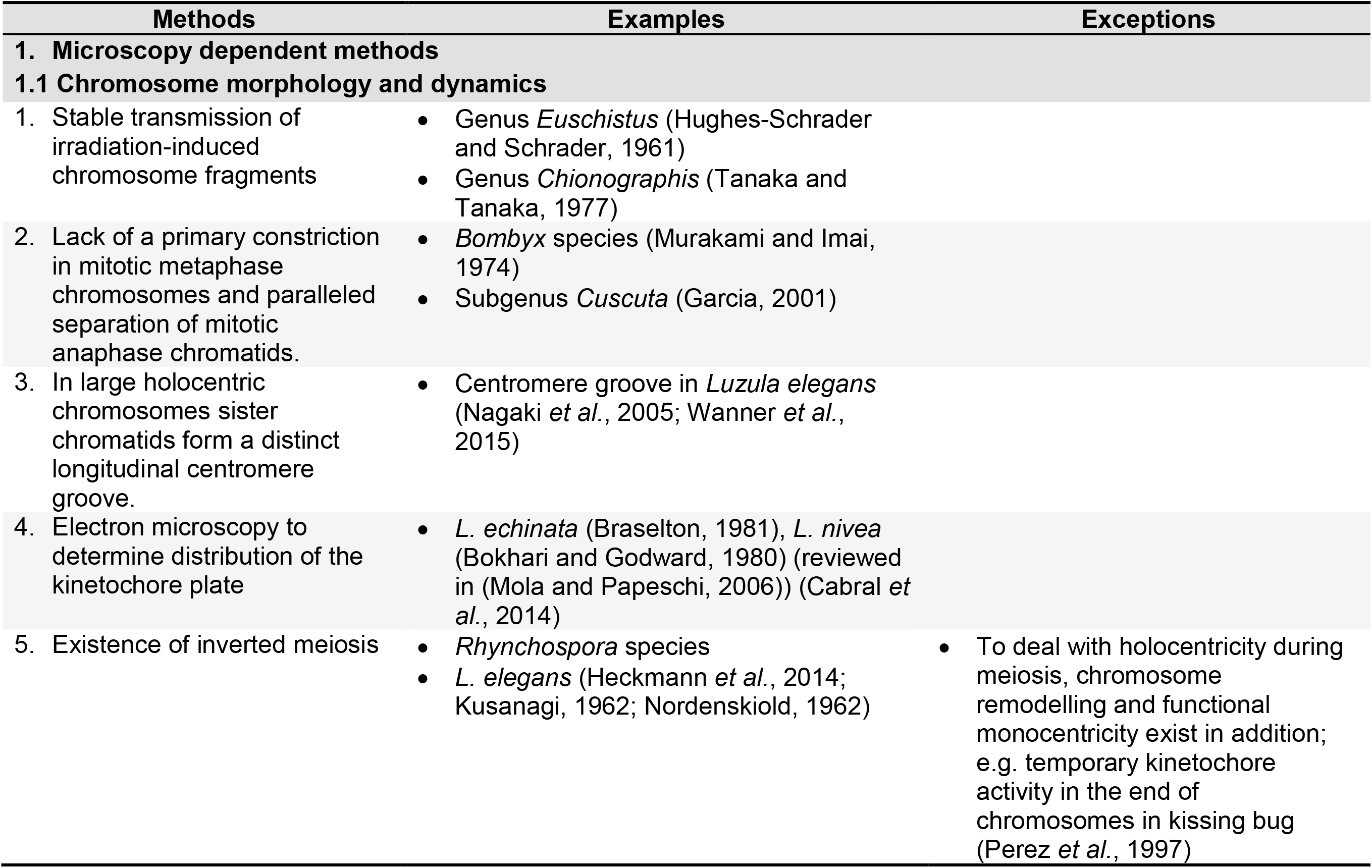
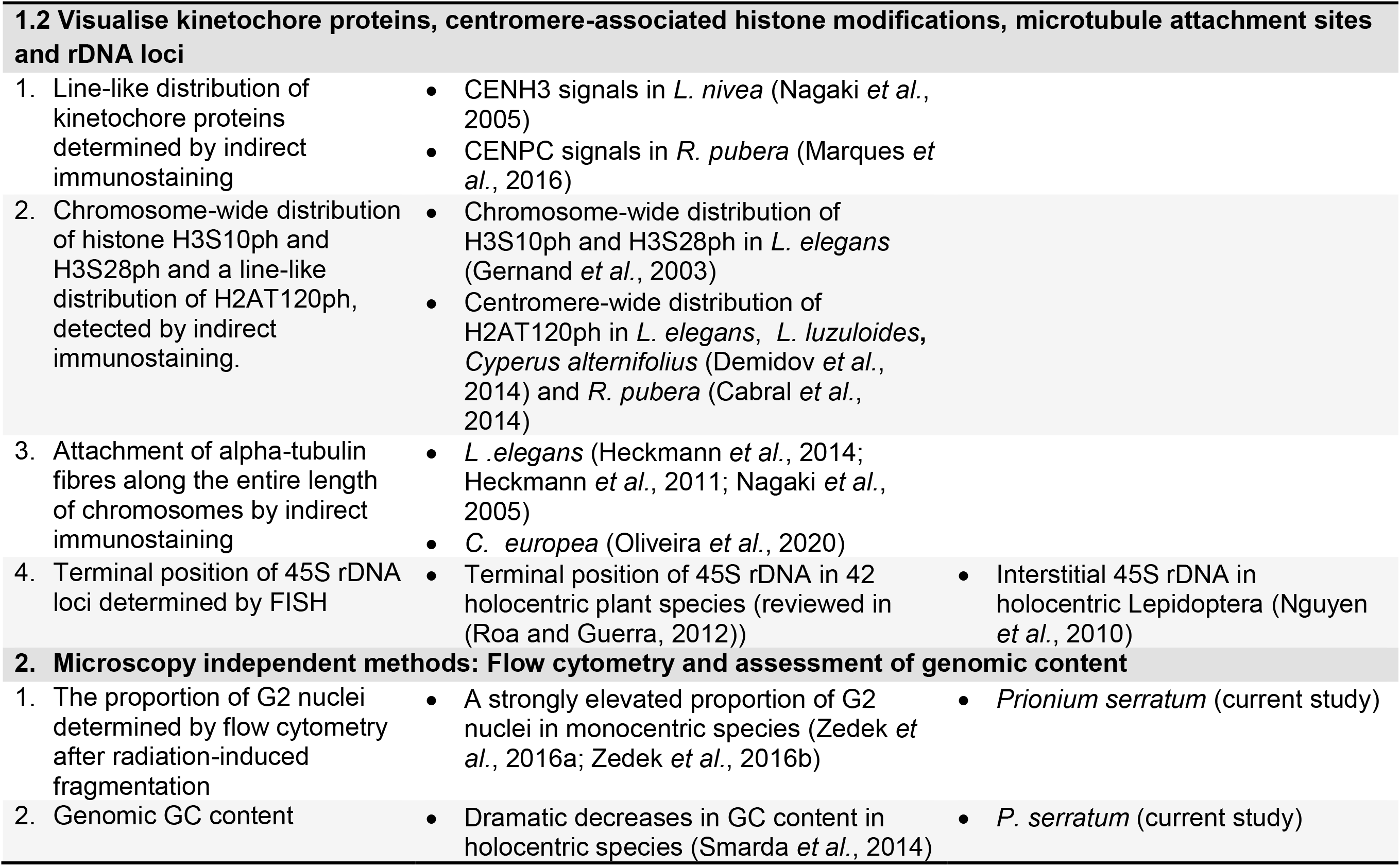
Methods to identify holocentricity

Cytological methods, by observing the absence of a primary constriction in mitotic chromosomes, paralleled segregation of anaphase sister chromatids and the faithful transmission of induced chromosomal fragments, are the prime methods of choice to identify holocentrics. In large chromosome species like *L. elegans* and *R. pubera*, holocentromeres form at somatic pro- and metaphase a distinct longitudinal groove along each sister chromatid which is visible by standard (Heckmann *et al.*, 2011; Nagaki *et al.*, 2005), structured illumination microscopy and scanning electron microscopy (Marques *et al.*, 2015; Wanner *et al.*, 2015).

However, the first two methods are not applicable for small chromosome species. The analysis of irradiation induced chromosome fragments is still one of the best methods to verify holocentricity (Hughes-Schrader and Ris, 1941; reviewed in Mola and Papeschi, (2006)). While acentric fragments of monocentric chromosomes form micronuclei, induced holocentric fragments are stably transmitted into the next cell generation and do not form micronuclei. But the application of this method requires specialised equipment for the generation of ionising radiation.

The analysis of meiotic chromosome dynamics has been used to determine holocentricity in species with moderate to large chromosomes (reviewed in ((Cuacos *et al.*, 2015); (Marques and Pedrosa-Harand, 2016)). Three principle options exist to deal with holocentricity during meiosis: (i) ‘chromosome remodelling’, (ii) ‘functional monocentricity’ and (iii) ‘inverted meiosis’. In the case of inverted meiosis, in contrast to monopolar sister centromere orientation, the unfused holokinetic sister centromeres behave as two distinct functional units during meiosis I, resulting in sister chromatid separation. Homologous non-sister chromatids remain terminally linked by a hardly visible chromatin fibre. Then, they separate at anaphase II. Thus, an inverted sequence of meiotic sister chromatid segregation occurs.

An almost terminal position of 45S rDNA, adjacent to telomeres, has been linked to holocentricity. This observation was made in 42 species of seven genera with holokinetic chromosomes (Roa and Guerra, 2012). A possible explanation is that a secondary constriction in the interstitial region would interrupt the kinetochore plate along the holokinetic chromosome establishing a condition similar to a dicentric chromosomes, leading to errors in chromosome segregation (Heckmann *et al.*, 2011). But in holocentric *Lepidoptera* species also interstitial 45S rDNA sites were detected (Nguyen *et al.*, 2010). Thus, since the terminal 45S rDNA location is not universal in holocentrics, it is not a universal evidence for holocentricity. Also, a terminal position of 45S rDNA was found in monocentric species (Schubert and Wobus, 1985).

Visualisation of kinetochore proteins, such as CENH3 or CENPC, by immunodetection shows the centromere type directly (Marques et al., 2016; Nagaki et al., 2005). This strategy is less restricted by chromosome size. However, it is often limited by the availability of species-specific kinetochore antibodies, which are both time- and cost-consuming in production. However, the absence of CENH3 in some species and the microtubule attachment at CENH3-free chromosome regions in some species (Drinnenberg *et al.*, 2014; Oliveira *et al.*, 2020) make the application of anti-CENH3 as a universal marker for centromeres questionable. Nevertheless, the analysis of kinetochore proteins could be complemented by combining the investigation of the spindle fibre attachment using alpha-tubulin-specific antibodies if the size of chromosomes allows the identification of the spindle fibre attachment site. In addition, the application of antibodies specific for the cell cycle-dependent pericentromeric phosphorylation of histone H3 (H3S10ph, H3S28ph) and H2A (H2AT120ph) resulted in the identification of holocentromere-specific immunostaining patterns (Demidov *et al.*, 2014; Gernand *et al.*, 2003). In monocentric plants, immunostaining with antibodies against H3S10ph and H3S28ph results in a specific labelling of the pericentromere in mitotic chromosomes. In contrast, in holocentric plants, immunolabelling with the same antibodies results in uniform staining of condensed chromosomes, due to the chromosome-wide distribution of the pericentromere (Gernand *et al.*, 2003). The application of these antibodies in a wide range of species is possible due to the evolutionarily conserved amino acid sequence of histone H3. However, in some monocentric species, the application of anti-H2AT120ph resulted in additional non-pericentromeric signals (Baez *et al.*, 2019; Sousa *et al.*, 2016).

Transmission electron microscopy studies also showed differences between holo- and monocentric chromosomes in relation to the size and distribution of the kinetochore plate (reviewed in (Mola and Papeschi, 2006)). However, the preparation of specimens for electron microscopy is somewhat laborious, and therefore it is less suitable for routine work.

Microscopy-independent flow cytometry and sequence-based approaches, by analysing irradiation-induced G2 nuclei accumulation and GC content, respectively, were developed for identifying the centromere type (Smarda *et al.*, 2014; Zedek *et al.*, 2016a). But, as our analysis of *P. serratum* showed, indirect methods should be taken with care. Hence, as no universal and straightforward method exists, if possible, different techniques should be combined to determine holocentricity, depending on the characteristics for each particular species.

## Acknowledgements

We would like to thank Tammy Elliot (University of Cape Town, South Africa), Boris O. Schlumpberger (Herrenhäuser Gardens, Germany) and Matthias Hoffmann (Botanical Garten Halle, Germany) for providing *P. serratum* plants. We thank Anne Fiebig (IPK) for die submission of sequence reads. This work has been supported by the Deutsche Forschungsgemeinschaft (HO1779/32-1), DAAD/CAPES (57517412; 88881.144086/2017-01) and Taiwan Ministry of Science and Technology (MOST, 106-2917-I-006-012 and 109-2917-I-564-022).

## Authors contributions

MB and YTK performed repeat analyses, FISH and immunostaining experiments and wrote the manuscript, YD performed FISH and immunostaining, TB determined the replication dynamics, AB *in silico* identified CENH3, JF measured the genome size, VS performed high resolution microscopy, ALLV, APH analysed data and AH designed the study, analysed data and wrote the manuscript.

## Supplementary Material

**Suppl. Figure 1.**
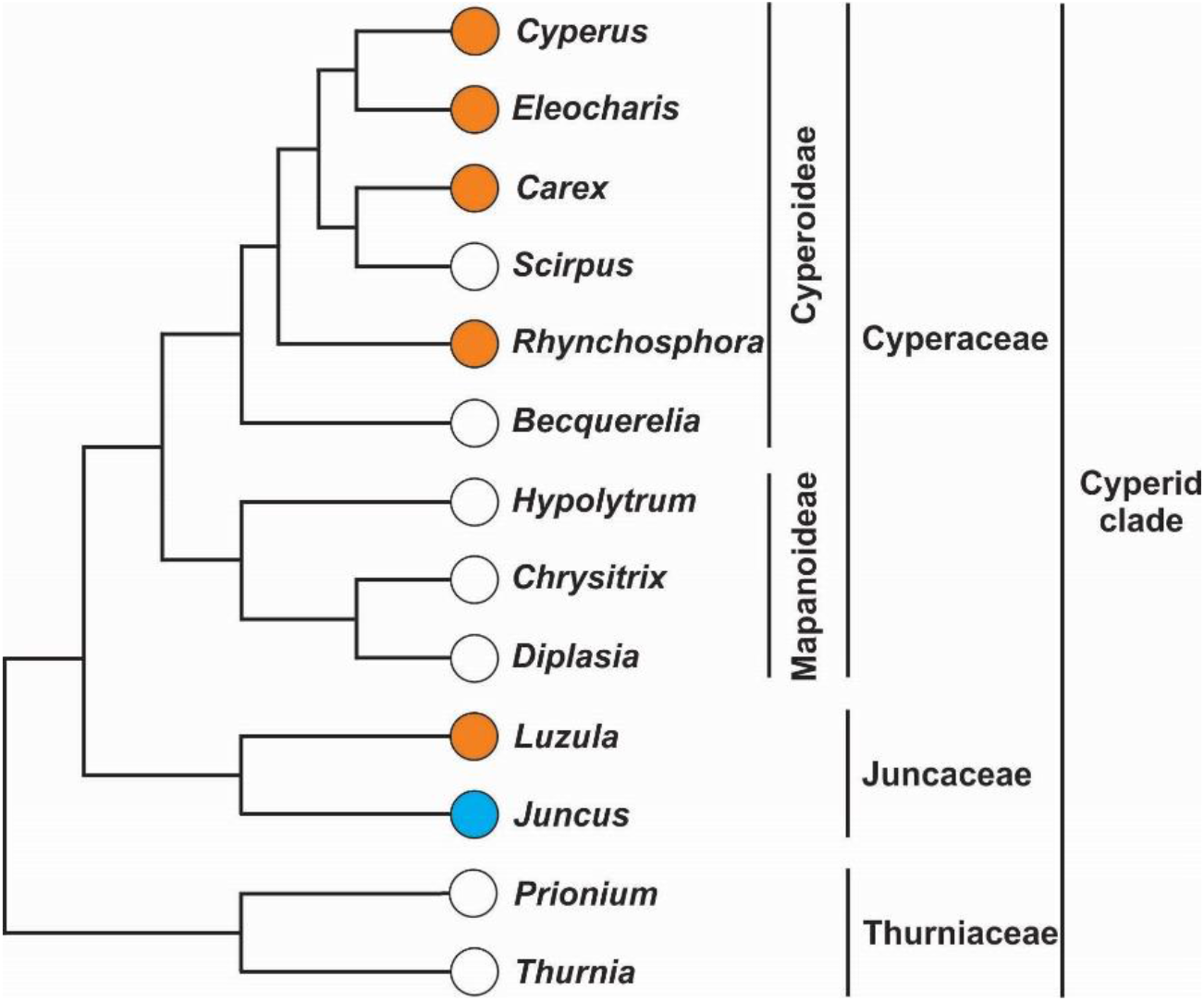
Phylogenetic relationship of the genera within the Cyperid clade. Confirmed holocentric genera are labelled by an orange circle; monocentric ones with a blue circle; genera with no centromere information with an empty circle. Phylogeny simplified from Hochbach *et al.* (2018), Semmouri *et al.* (2019), Silva *et al.*, (2020).

